# A genome engineering resource to uncover principles of cellular organization and tissue architecture by lipid signalling

**DOI:** 10.1101/2020.02.03.933093

**Authors:** Deepti Trivedi, Vinitha CM, Karishma Bisht, Vishnu Janardan, Awadhesh Pandit, Bishal Basak, Padinjat Raghu

## Abstract

Phosphoinositides (PI) are key regulators of cellular organization in eukaryotes and genes that tune PI signalling are implicated in human disease mechanisms. Biochemical analyses and studies in cultured cells have identified a large number of proteins that can mediate PI signalling. However, the role of such proteins in regulating cellular processes *in vivo* and development in metazoans remains to be understood. Here we describe a set of CRISPR based genome engineering tools that allow the manipulation of each of these proteins with spatial and temporal control during metazoan development. We demonstrate the use of these reagents to deplete a set of 103 proteins individually in the *Drosophila* eye and identify several new molecules that control eye development. Our work demonstrates the power of this resource in uncovering the molecular basis of tissue homeostasis during normal development and in human disease biology.

## Introduction

Phosphoinositides (PI) are low abundance phospholipids implicated in the regulation of many important biological processes including cell polarity, cell migration, aging, growth and development (Balla, 2013). The head group of the parent lipid phosphatidylinositol can be phosphorylated combinatorially at positions 3, 4 and 5 to generate a set of seven PIs. These PIs are key regulators of sub-cellular processes; for e.g., phosphatidylinositol 4,5 bisphosphate [PI(4,5)P_2_] plays a key role in the regulation of membrane transport and the cytoskeleton (Janmey et al., 2018) while phosphatidylinositol 3 phosphate [PI3P] is an important regulator of endocytosis and autophagy (Schink et al., 2016). PIs have also been implicated in the regulation of nuclear function (Fiume et al., 2015). PIs also play a key role in regulating developmental processes in metazoans. For example, phosphatidylinositol 3,4,5 trisphosphate (PIP_3_) is an essential regulator of growth factor signalling and plays a conserved role in developmental control in *C. elegans, Drosophila* and mammals. Phosphatidylinositol 4-phosphate (PI4P) levels play a key role during cellular development in gametogenesis in *Drosophila* (Brill et al., 2000; Tan et al., 2014) and a conserved role for PI(4,5)P_2_ is proposed in apico-basal polarity (Devergne et al., 2014; Martin-Belmonte et al., 2007; Rousso et al., 2013; Yan et al., 2011). Dysregulation in PI signalling has also been linked to several human diseases; these include developmental disorders such as Lowe syndrome (Staiano et al., 2015) and PIK3CA-related overgrowth spectrum (PROS) (Madsen et al., 2018); mutations in PTEN, a key regulator of PIP_3_ levels are the second most frequent mutations seen in human cancers (Worby and Dixon, 2014). Mutations in genes regulating PI signalling are seen in a large number of genetic disorders of the human nervous system (Raghu et al., 2019). Thus, the PI signalling pathway is a key regulator of cell function and tissue architecture both during normal development and also in disease mechanisms and understanding their role in such processes is of fundamental importance.

Given their importance in a large number of important sub-cellular processes, PI signalling is tightly regulated. The seven PIs are generated in cells by the activity of a set of evolutionarily conserved lipid kinases that are specific for the substrate they will use as well as the position on the inositol at which they will phosphorylate (Sasaki et al., 2009). In turn, once generated, PIs exert their cellular effects by binding to and modulating the activity of a large number of PI binding proteins whose functions are key to the control of sub-cellular processes by these lipids. For example many proteins contain domains that bind PI3P (PX and FYVE) and PI4P (FAPP and OSBP) and numerous PH domains that bind various PIs with differing degrees of affinity and selectivity have been reported (Hammond and Balla, 2015). Indeed, a recent biochemical study has identified a large number of proteins that bind a range of PIs with varying degrees of specificity (Jungmichel et al., 2014). However, the biological function of many of these proteins remains to be discovered.

Although PI signalling is broadly a conserved feature of all eukaryotic cells, some aspects of this pathway are found uniquely in metazoa and play important roles in animal development and tissue architecture. However, there remain key challenges in the analysis of PI signalling in metazoan development. For example, PIs may regulate the same sub-cellular process (e.g endocytosis) but may be co-opted to control the trafficking of unique cargoes in different tissues. Second, through their ability to control the secretion of intercellular signalling morphogens and hormones that mediate intercellular signalling, PIs can regulate tissue development in a non-cell autonomous manner. Third, given their key functions in cell biology, whole body knockouts of these genes often result in organismal lethality. Finally, in many cases, a given biochemical activity is underpinned by multiple genes in mammalian genomes and gene redundancy usually makes analysis of *in vivo* function challenging.

Analysis in *Drosophila* offers a powerful alternative to overcome the challenges of studying PI function in cell and developmental processes. Many developmental mechanisms are conserved between *Drosophila* and other metazoan models. A detailed annotation of the PI signalling genes in the *Drosophila* genome (Balakrishnan et al., 2015) led to the identification of 103 genes which are part of the PI signalling toolkit (Figure 1) and conserved between flies and other organisms important for the study of developmental processes and disease biology. Further the availability of binary gene expression control systems such as the GAL4/UAS system allow for tight spatial and temporal gene modulation in turn allowing sophisticated analysis of both cell autonomous and non-cell autonomous modes of developmental control by PI signalling (Colombani et al., 2005). Thus, *Drosophila* offers a model where one can analyze the control of cellular and developmental processes by PI signalling in a metazoan context.

**Figure 1:**
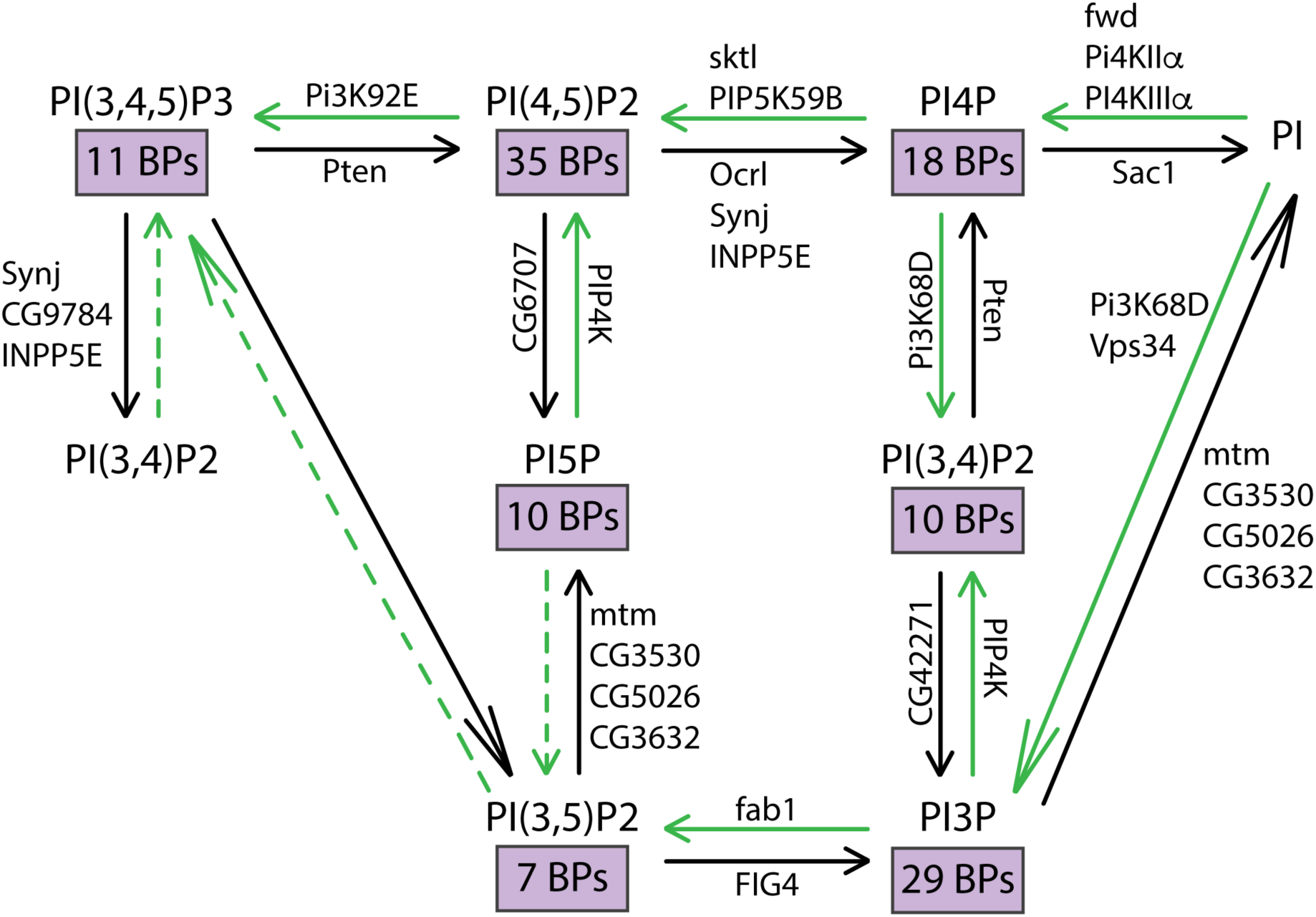
Interconversion of phosphoinositides in eukaryotic cells. The metabolic pathways by which the different phosphorylated forms of phosphatidylinositol are interconverted are represented. The kinase and phosphatase reactions are indicated by green and black arrows respectively. The *Drosophila* gene encoding each of the enzymes responsible for these reactions are also indicated. Dotted lines indicate reactions that are yet to be established. The number of binding proteins (BPs) capable of binding each of the phosphoinositides are indicated in purple boxes.

In the recent years, CRISPR/Cas9 has emerged as a powerful tool for genome editing in multiple organisms including flies (Gratz et al., 2013; Wang et al., 2016; Wu et al., 2018). However, unlike transgenic fly lines expressing RNAi constructs that are available against almost the entire fly genome (Dietzl et al., 2007; Perkins et al., 2015), efforts to make guide RNA (gRNA) expressing transgenic flies are still underway (Meltzer et al., 2019; Port et al., 2019). In this study, we have generated a genome editing system that can be used for Cas9 mediated editing to delete the open reading frame of each of the 103 annotated genes that could regulate PI signalling (Balakrishnan et al., 2015). We demonstrate the use of this resource both in cultured *Drosophila* cells as well as in intact *Drosophila* tissues *in vivo* during development. Importantly, we use these guides to generate both whole body knock outs as well as tissue specific, developmentally timed depletion of individual genes with high efficiency. Further, using these reagents we present the results of a genetic screen in the developing *Drosophila* eye and identify key roles for PI signalling in this influential model of tissue development and patterning. Our reagent set will be a powerful and versatile tool for the discovery of novel regulation by PI signalling of development, tissue homeostasis and human disease biology.

## Results

### Design of gRNAs for editing PI signalling genes

In designing the gRNAs to target each PI signalling gene, we considered a few important factors: Firstly, CRISPR-Cas9 based genome editing is highly robust and specific; a single mismatch in the target sequence may highly reduce the efficiency of CRISPR (Pattanayak et al., 2013), especially if the mismatches are present in the core 12bp protospacer region most proximal to the protospacer adjacent motif (PAM) site. Second, a large number of users may find the library of dgRNA transgenic flies useful to study PI signalling in their own research contexts. This would mean that the dgRNA lines may be used in flies of different genetic backgrounds. Third, the same dgRNA constructs should be able to target the genes in S2R+ cells for editing prior to biochemical experiments. Considering these points, we decided not to directly use the reference genome sequence to design the gRNAs. The entire genome of S2R+ cells as well as the isogenised parent fly stock (BL# 25709) that was to be used to generate dgRNA transgenic flies were sequenced (Sequencing data is available on hyperlink). Sequences thus obtained were aligned against the reference genome and also against each other. When aligned against the reference genome, both the isogenised parent fly stock and the S2R+ cell genome sequences showed more than 600,000 single nucleotide polymorphisms (SNPs). When compared to each other, these genome sequences had ∼1.3 million mismatches (SNP, insertions and deletions) suggesting that, as expected, almost all of the sequence differences in the 2 genomes were independent of each other (Figure 2). While designing the single gRNA (sgRNA), we only chose those targets that did not have sequence mismatches in any of the 3 genomes (reference genome and the 2 genomes sequenced) thus ensuring that the dgRNAs could be used in flies of diverse genetic backgrounds and also in S2R+ cells. In addition, this allowed for a relatively simple way to test the designed dgRNAs in S2R+ cells for their ability to target specific genes prior to generating transgenic fly lines.

**Figure 2:**
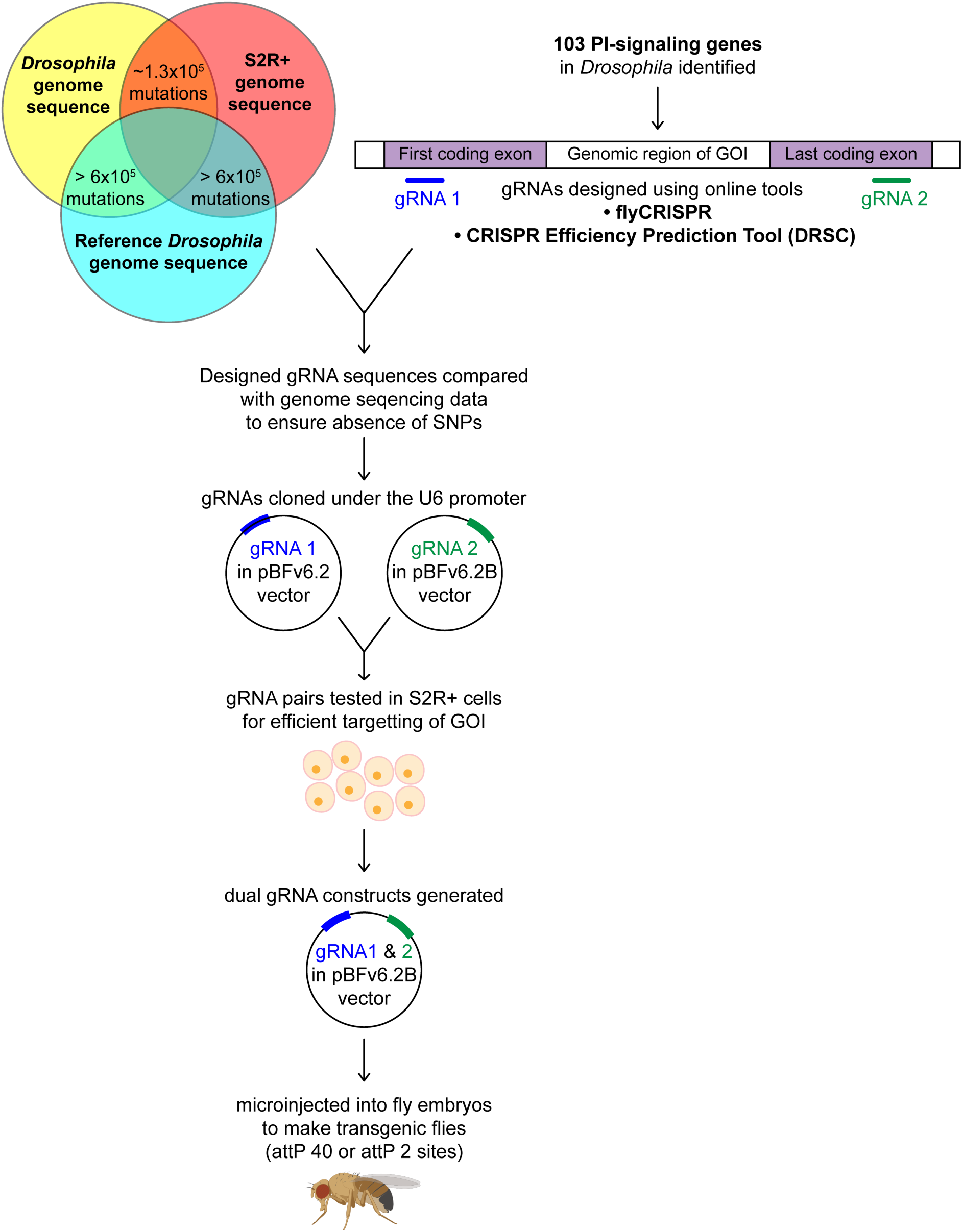
Workflow for the generation of dgRNA transgenic flies. For each of the 103 PI-signalling genes in the *Drosophila* genome, two gRNAs were designed using flyCRISPR and verified using the CRISPR Efficiency Prediction Tool (DRSC). The first gRNA (gRNA 1; indicated in blue) was designed to target the first coding exon and the second gRNA (gRNA 2; indicated in green) was designed to target the last coding exon. It was ensured that these designed gRNAs did not have any mismatches when compared against any of the 3 genomes (the genomes of *Drosophila* BL25709 line used for microinjection, S2R+ cells and the reference genome). The two gRNAs (gRNA 1 and gRNA 2) for each gene were cloned into pBFv6.2 and pBFv6.2B respectively and tested in S2R+ cells for their ability to delete the target genes in the presence of Cas9. Following this, both the gRNAs for each gene were cloned into a single plasmid to generate the dgRNA constructs. These were microinjected into *Drosophila* embryos to generate dgRNA transgenic flies against each of the PI-signalling genes.

For each gene, two regions were identified wherein a sgRNA would be designed to target each region. The first sgRNA would be designed to target the first coding exon and the second sgRNA to target the exon with the stop codon (Figure 2). In case of genes with alternate spliced forms, regions around the most 5’ start codon and the most 3’ stop codon were chosen (For example, CG3682 has upto 6 splice variants with 4 putative start sites). In cases where two or more putative genes are annotated on the same locus, sgRNA target regions were chosen such that the coding sequences of neighbouring genes are not disrupted (For example, there are 6 different putative ORFs in the first few introns of *plc21c* and hence the first sgRNA is designed on exon 8). Once a set of two ∼100 bp regions were identified for each gene keeping in mind the above-mentioned criteria, we designed the sgRNAs using an online tool (http://targetfinder.flycrispr.neuro.brown.edu) with zero predicted off-targets. During the course of this study, an additional online tool (https://www.flyrnai.org/evaluateCrispr/) became available. Thereafter, this tool was used to obtain a predicted efficiency score. The most efficient sgRNAs in the region were then checked in our sequence database for presence of any mismatches in either S2R+ cells or injection fly stocks compared to the reference genome sequence. The best sgRNA sequences predicted to efficiently target specific genes with no predicted off-targets and absence of any mismatches were chosen for synthesis. Any sgRNA sequences that did not qualify these criteria were discarded and new sgRNAs designed to fit all of the above criteria.

### Generation of gRNA expression constructs

The first sgRNAs targeting the start codon for each gene were cloned into the pBFv6.2 vector and the second sgRNAs targeting the stop codon into the pBFv6.2B vector. Both of these vectors ubiquitously express the gRNAs under the U6.2 promoter (Kondo and Ueda, 2013). The two sgRNA constructs targeting a given gene along with pUAST-Cas9-T2A-eGFP were co-transfected into S2R+ cells stably expressing Tubulin-Gal4 (Gift from Satyajit Mayor). The sgRNA pairs that led to the deletion of the gene, as tested by PCR and sequencing of the target gene loci, were chosen for dual-gRNA (dgRNA) construction. In case a particular pair of sgRNAs failed to delete the target gene, a new set of sgRNAs were designed for both the first and the last coding exon and different combinations of sgRNAs were tested until the most optimal pair capable of deleting the target gene was identified.

Once a functional pair of sgRNAs capable of deleting a target gene was identified, the first sgRNA was cloned into the pBFv6.2B vector containing the second sgRNA to generate the dgRNA construct (*U6-dgRNA*). These constructs were then microinjected to generate dgRNA transgenic flies for each of the 103 PI signalling genes (Figure 2). This U6-dgRNA construct can also be transfected in S2R+ cells to generate gene deletions for cell culture-based experiments. For genes that are present on the 2^nd^ chromosome, the dgRNA construct was inserted on the 3^rd^ chromosome (Table I). For all the other genes, the dgRNA was inserted on the 2^nd^ chromosome (See materials and methods). This design offers the advantage that when used to generate whole fly knockouts, the dgRNA transgenes can subsequently be readily out-crossed given that the gRNA and the target gene are on two different chromosomes.

**Table I:**
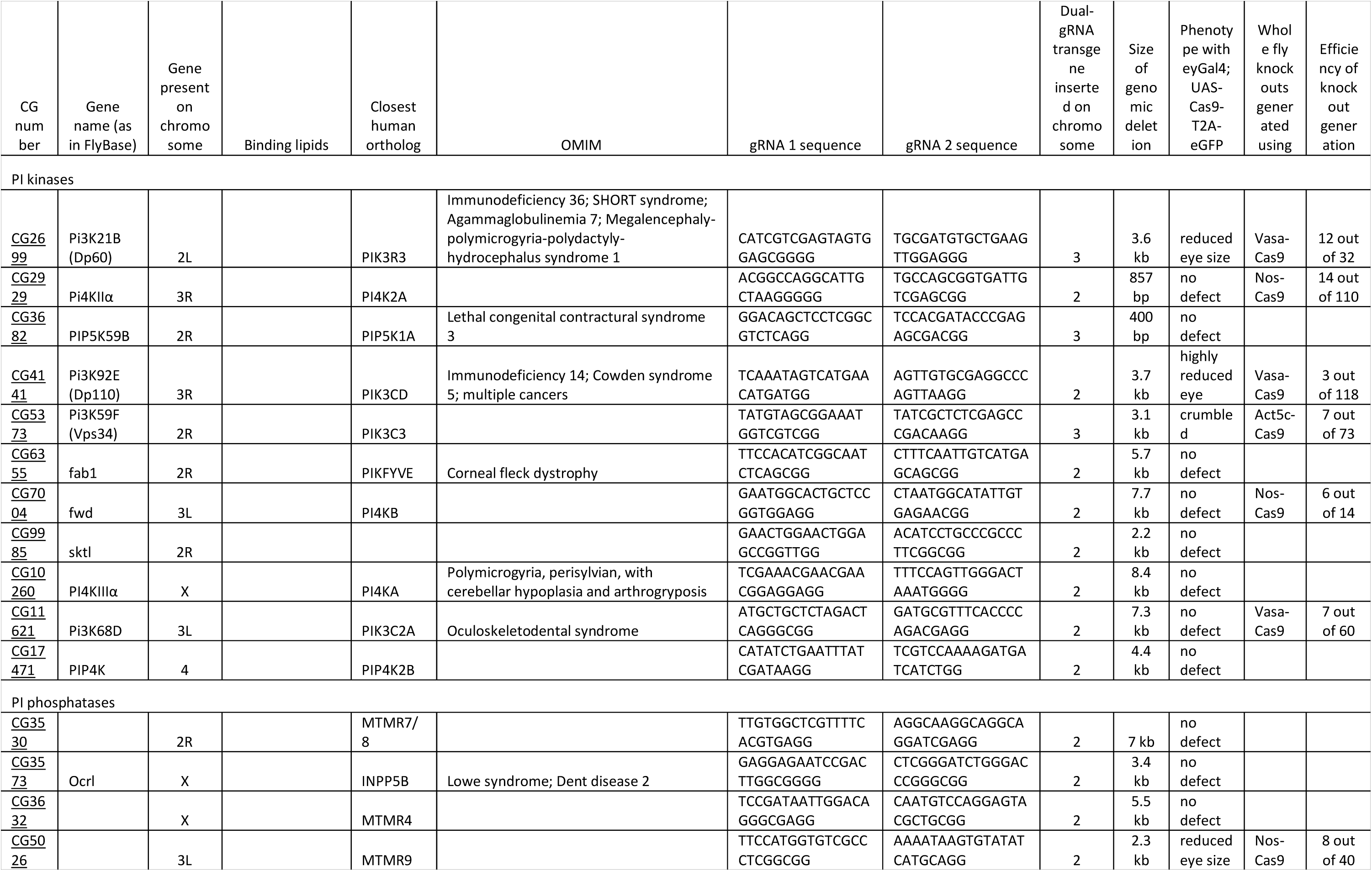
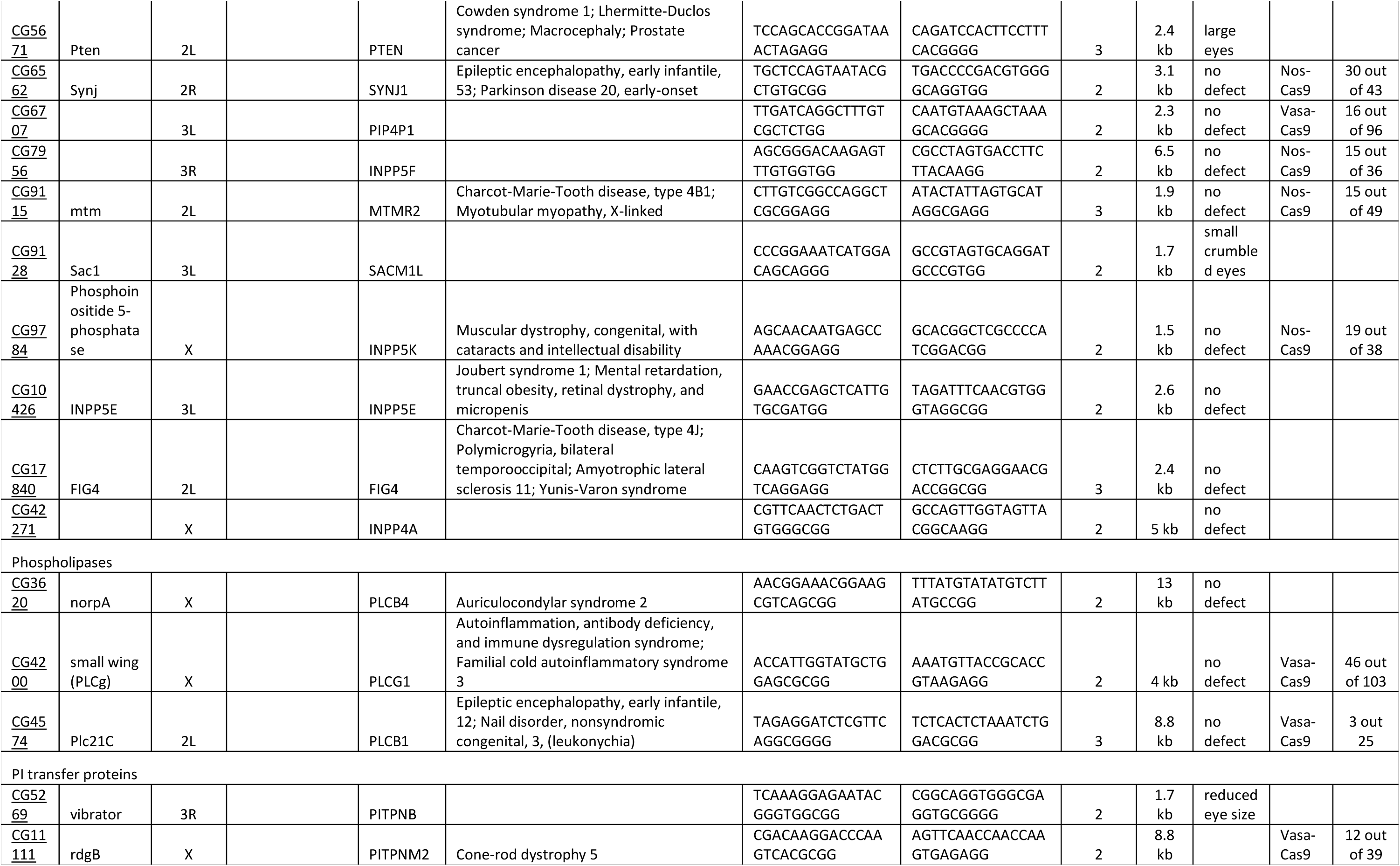
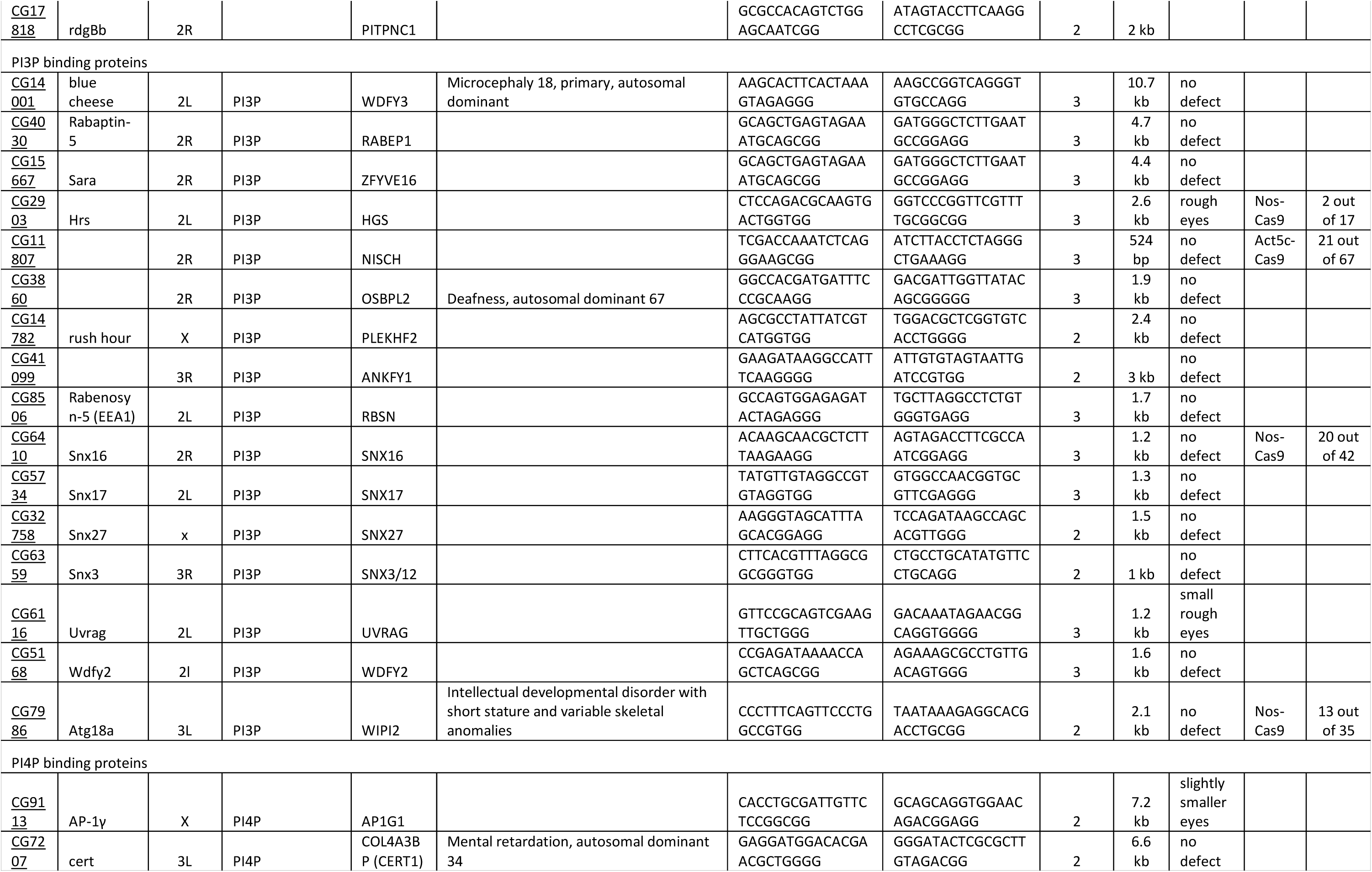
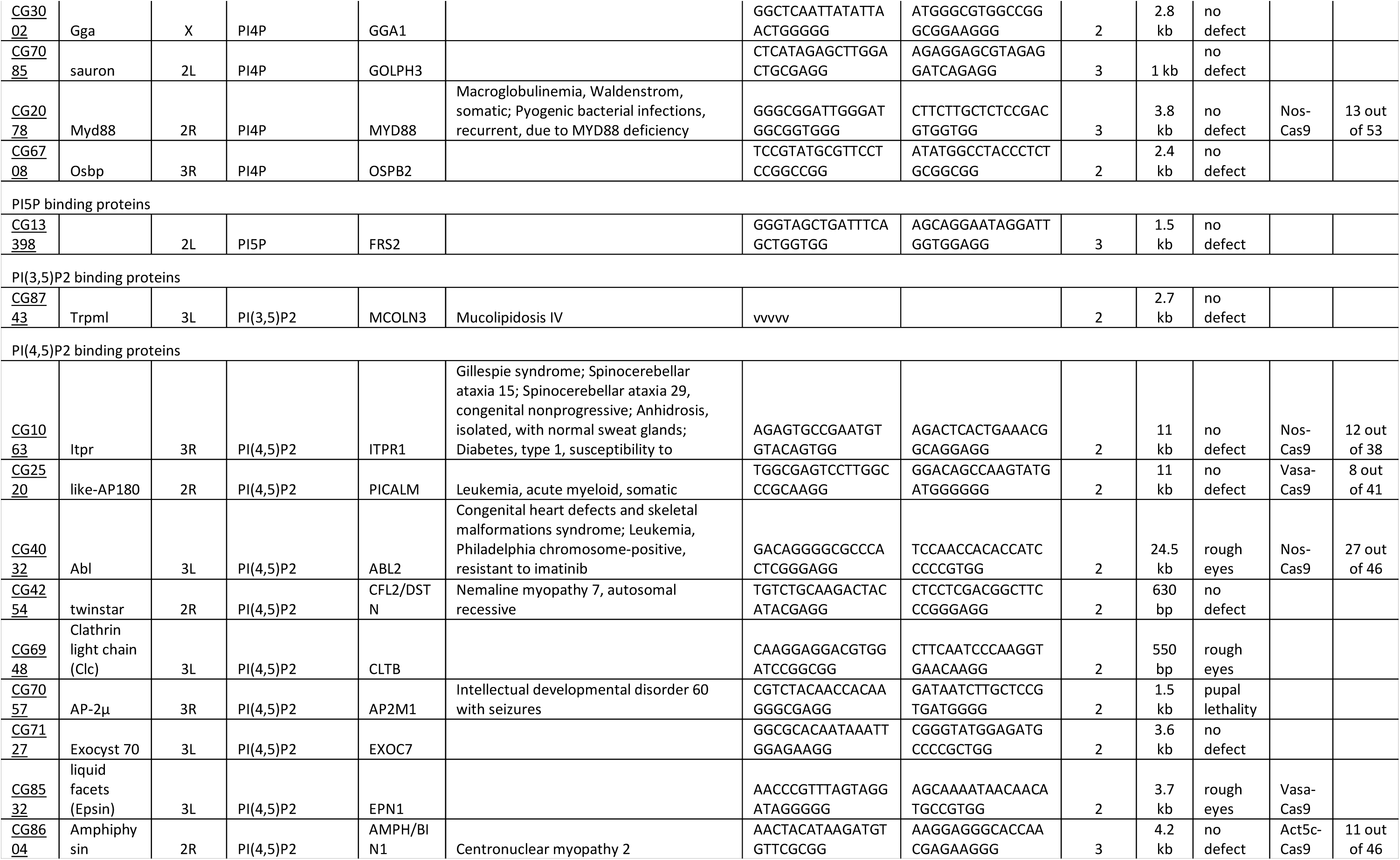
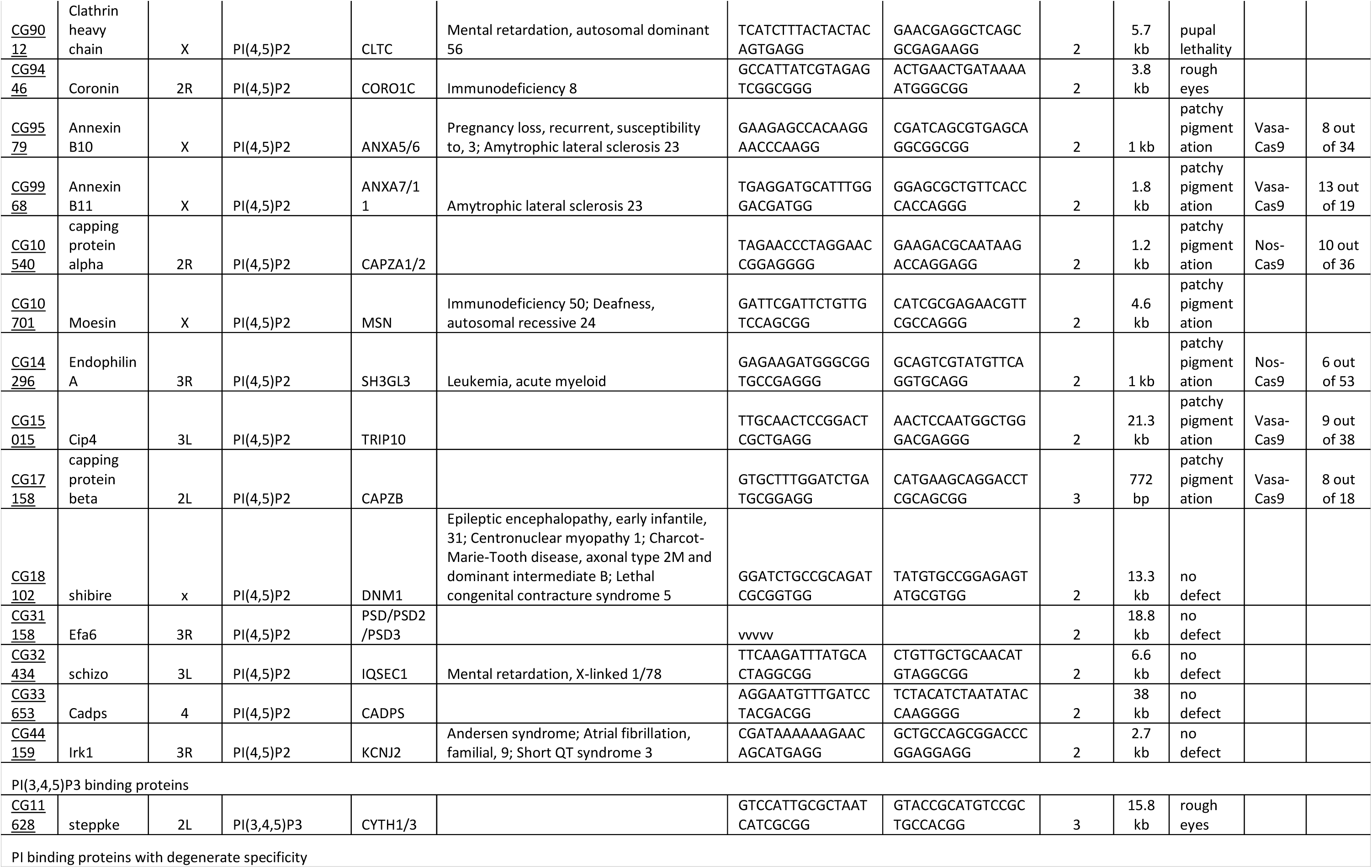
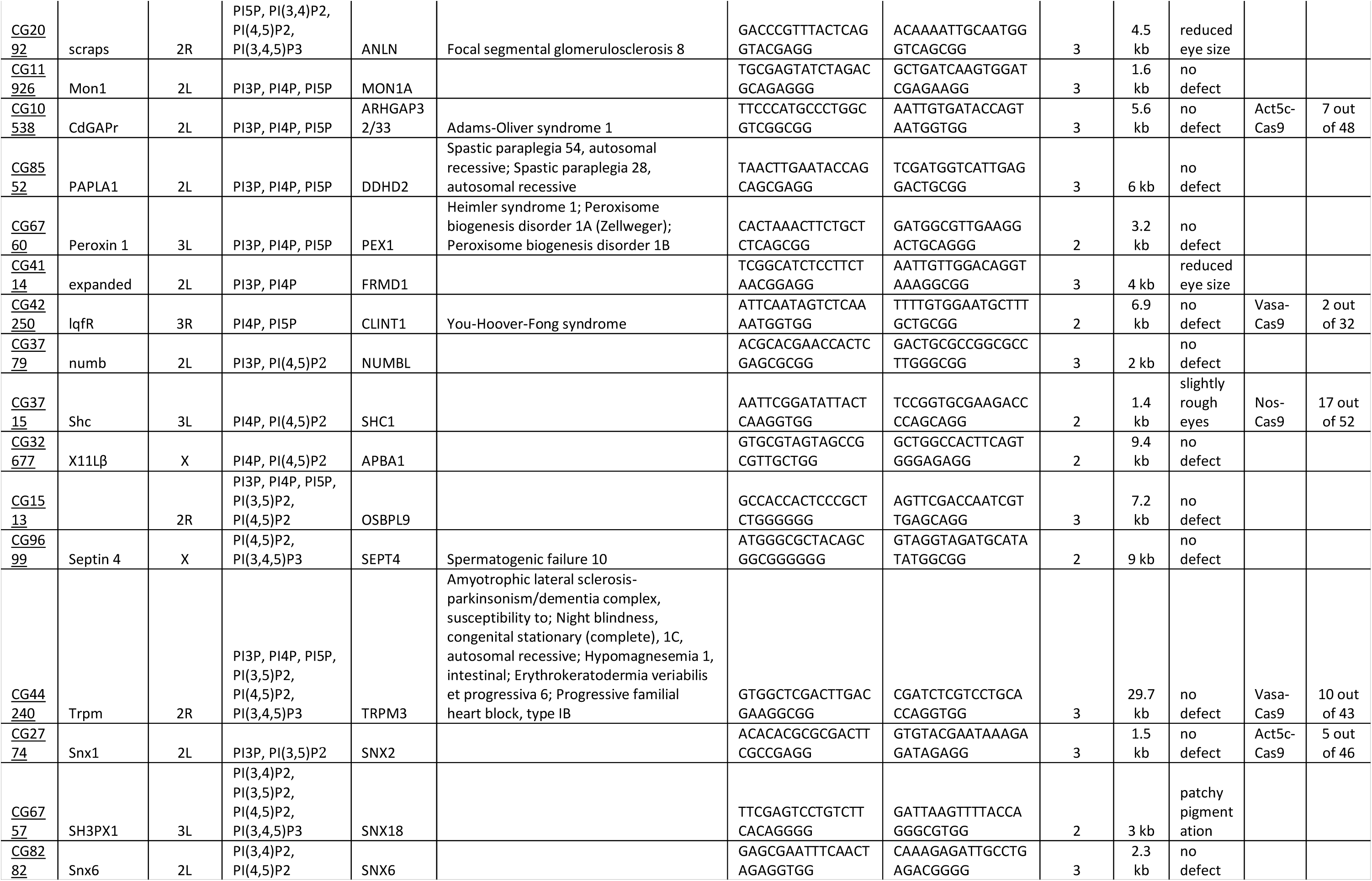
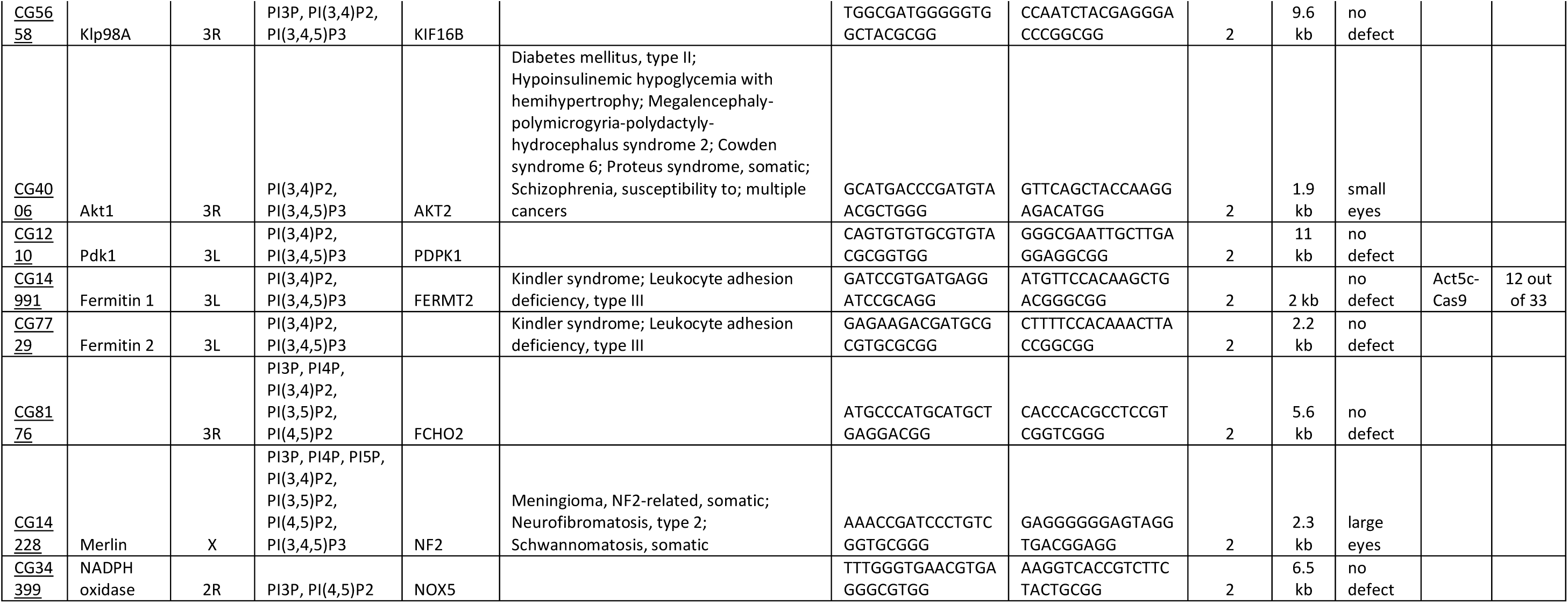
List of all 103 *Drosophila* PI-signalling genes against which dgRNAs have been generated. The table indicates the CG numbers, gene names and what chromosome each of the genes are located on. For the phosphoinositide binding proteins, the various phosphoinositides either established or predicted to bind each protein have been listed. The table also has the closest human orthologs and associated human diseases from the Online Mendelian Inheritance in Man (OMIM) database indicated. The sequences of gRNA 1 and gRNA 2 for each gene are listed along with the size of the genomic deletion expected from these gRNA combinations, the phenotypes obtained when these genes were deleted specifically in the eye and the efficiency of whole fly gene knockout generation using either Vasa-Cas9, Act5c-Cas9 or Nos-Cas9.

### Generation of UAS-Cas9-eGFP transgenic lines

At the core of CRISPR/Cas9 technology is the endonuclease Cas9 which utilizes gRNAs to target specific genomic sequences and generate a double stranded break. Transgenic flies expressing Cas9 under UAS control are already available (Port et al., 2019). However, these lack a reporter and therefore it is difficult to readily identify cells or tissues expressing Cas9, which would be of great advantage when targeting genes in a tissue specific manner. In order to satisfy this requirement, we designed and generated transgenic UAS-Cas9 flies with a fluorescence reporter eGFP. Given that the human codon optimised Cas9 (Cas9.P2) expresses at lower levels thereby reducing the cytotoxic effects of the otherwise highly expressing Cas9.C (Meltzer et al., 2019), we have used Cas9.P2 to generate our construct. In order to ensure that the presence of an eGFP tag does not hinder the activity of Cas9, we introduced a self-cleaving peptide T2A sequence in between Cas9 and eGFP (Figure 3A). This ensured that while the Cas9 and eGFP are expressed from the same mRNA, they are made as two independent proteins. Presence of the eGFP coding sequence downstream of Cas9 ensured that every cell positive for eGFP fluorescence would definitely also express Cas9. In addition, we tagged the Cas9 with a nuclear localization signal (NLS) at both the N- and the C-terminal to facilitate its translocation into the nucleus for better access of the genomic DNA.

**Figure 3:**
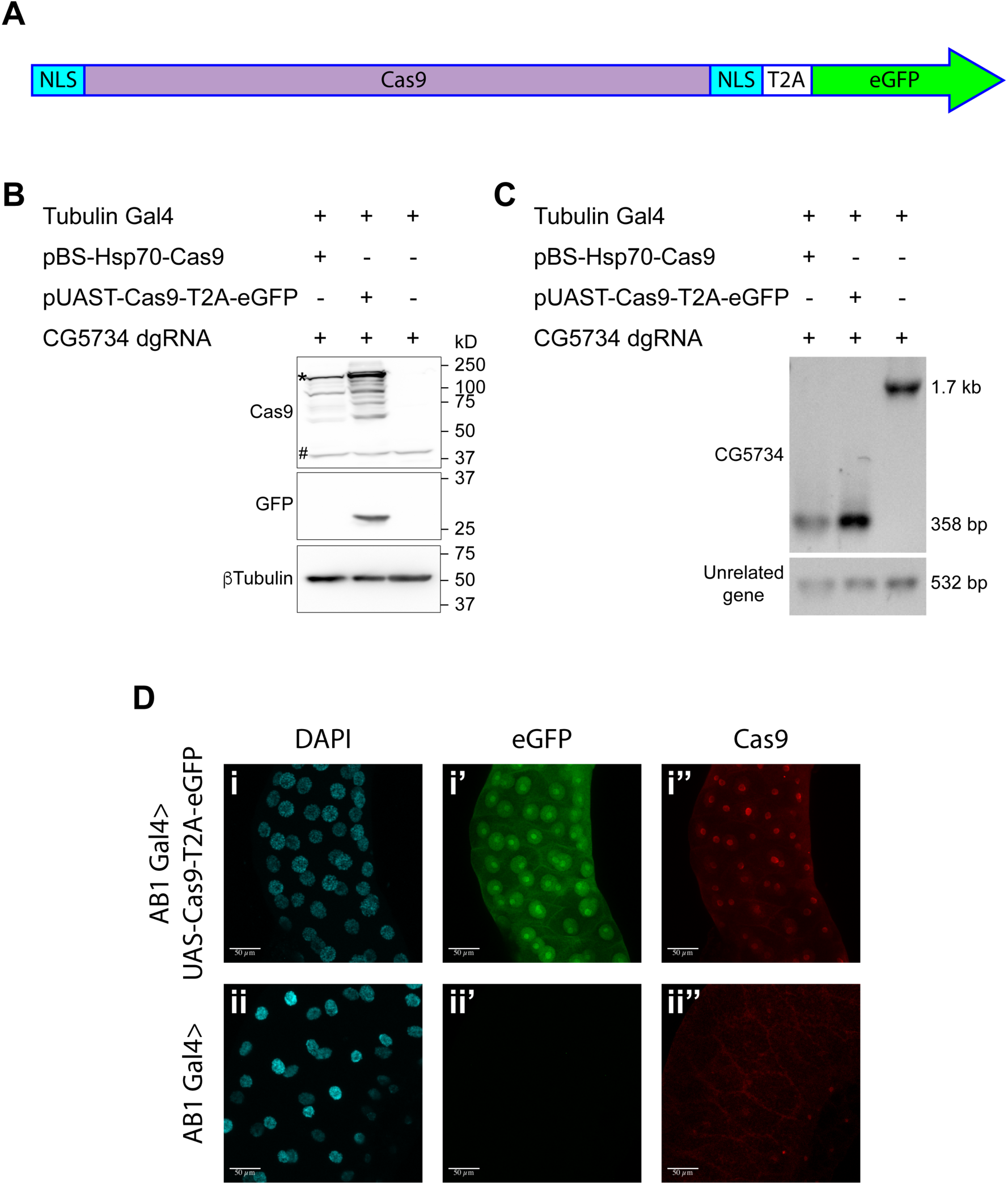
Design and validation of UAS-Cas9-T2A-eGFP. (A) A schematic of the Cas9-T2A-eGFP construct indicating the presence of a nuclear localization signal (NLS) at the N and C termini of Cas9. An eGFP sequence is present downstream of the Cas9 sequence and these two are separated by the T2A sequence. (B) Western blot of S2R+ cells expressing Cas9-T2A-eGFP. S2R+ cells constitutively expressing Tubulin Gal4 were transiently transfected with the indicated plasmids. Cells were harvested 48h after transfection and one half of the cells were subjected to Western blot analysis to verify that the T2A sequence was efficient and therefore resulted in expression of Cas9 and eGFP as independent proteins. The expected molecular size of Cas9 is 158 kD (the band of highest intensity indicated with a ‘*’). Note that the Cas9 antibody cross-reacts with a protein from S2R+ cells (‘#’) thereby precluding immunocytochemical detection of Cas9 in S2R+ cells. (C) Genomic PCR was performed on the other half of the cells to test for the ability of Cas9 to target CG5734. The presence of a 358 bp band in cells expressing Cas9 compared to the 1.7 kb band in untransfected cells suggests the successful deletion of CG5734. (D) UAS-Cas9-T2A-eGFP transgenic flies generated were crossed to AB1 Gal4 flies and the salivary glands of the progeny dissected and stained for Cas9. Cas9 was predominantly localized in the nucleolus suggesting efficient nuclear localization by the NLSs. Cells expressing Cas9 were marked by eGFP. Scale bar is 50 µm.

The UAS-Cas9-T2A-eGFP construct thus generated was tested for its genome editing efficiency and the usefulness of eGFP as a reporter. To test the Cas9, the pUAST-Cas9-T2A-eGFP construct was transfected along with the dgRNA against CG5734 (a gene predicted to have a PH domain that may bind to PI3P) into S2R+ cells constitutively expressing Tubulin-Gal4. As a control, the cells were parallelly co-transfected with the pBS-hsp70-Cas9 plasmid (Addgene Plasmid #46294) and the dual-gRNA against CG5734. Forty-eight hours post transfection the cells were harvested. Half the cells were used for protein extraction and Western blotting. Western blotting showed that Cas9 was being expressed from the UAS-Cas9-T2A-eGFP construct. The molecular size of Cas9 from this construct was similar to Cas9 expressed from pBS-hsp70-Cas9. Detection of a band corresponding to the molecular weight of free eGFP in cells transfected with UAS-Cas9-T2A-eGFP suggest that the T2A sequence was working efficiently to generate Cas9 and eGFP as two independent proteins from a single mRNA (Figure 3B). From the other half of the cells, genomic DNA was isolated and PCR performed to detect deletions in CG5734. We found the predicted deletion fragment generated by both the Cas9 from the newly generated pUAST-Cas9-T2A-eGFP plasmid and the Cas9 expressed from the pBS-hsp70-Cas9 control plasmid (Figure 3C) thus verifying that Cas9 expressed from the pUAST-Cas9-T2A-eGFP construct was capable of deleting target genes in the presence of appropriate gRNAs.

After verification in S2R+ cells, the pUAST-Cas9-T2A-eGFP construct was microinjected into fly embryos along with a helper plasmid expressing transposase to facilitate random P-element based insertion and obtain transgenic flies. In order to verify that the fly line obtained expresses Cas9, we crossed these lines to salivary gland specific Gal4 (AB1Gal4) flies. The progeny flies were dissected, salivary glands stained for Cas9 and imaged. All salivary gland cells expressed eGFP and were also stained positive for Cas9 thus demonstrating that the pUAST-Cas9-T2A-eGFP construct can be used as expected to drive Cas9 expression and to mark the Cas9 expressing cells with eGFP (Figure 3D). The eGFP expression was predominantly nuclear although some cytosolic expression could be observed. Cas9 did not have the same intracellular localization thus suggesting that the T2A sequence was functional and that the two proteins, Cas9 and eGFP were being expressed as independent proteins. Owing to the NLSs attached to the Cas9, its expression was limited to the nucleus. The pUAST-Cas9-T2A-eGFP flies showed Cas9 expression similar to the already existing Cas9 transgenic flies (Bloom number 54592) with the added advantage of eGFP expression to mark Cas9 expressing cells. These pUAST-Cas9-T2A-eGFP flies were used for all experiments described further.

### Investigating the role of PI signalling in eye development using the dgRNA toolkit

The *Drosophila* eye serves as an excellent model to perform genetic screens as it is not essential for viability and offers a phenotype that can be easily scored. Previously, influential studies have used *Drosophila* eyes to screen for genes regulating cell growth and proliferation [reviewed in(Tseng and Hariharan, 2002)]. The *Drosophila* eye is composed of ∼800 regularly arranged ommatidia, each containing 8 photoreceptor neurons making the *Drosophila* eye an attractive tissue to study neurodegenerative diseases like Huntington’s disease (Krench and Littleton, 2013) and screen for genes and genetic modifiers involved in neural development (Ma et al., 2009). Moreover, photoreceptor neurons are an excellent model for phototransduction which in flies is largely dependent on phosphoinositide signalling (Raghu et al., 2012) and dysregulation of PI signalling leads to defects in the development, cellular structure and functions of the eye.

To test the use of our dgRNA transgenic library, we studied the effect of deletion of PI signalling genes in the eye disc during development by expressing Cas9-T2A-eGFP using eyeless-Gal4 (eyGal4). *eyeless (ey)* is expressed very early on in the eye primordia in the embryo and before the formation of the morphogenetic furrow at the time of photoreceptor determination in the third instar larva (Halder et al., 1995). In addition, *ey* is also expressed in neurons. To delete target genes in a tissue specific manner we generated *ey-Gal4; UAS-Cas9-T2A-eGFP* flies and crossed them individually to each of the 103 *U6-dgRNA* transgenic lines. Prior to analysing the phenotypes obtained in the progeny of these crosses, we tested whether the gene under study was being edited in a tissue specific manner. For this, we isolated genomic DNA from the head and the body of a few representative *ey*>*GAL4; UAS-Cas9-T2A-eGFP; U6-dgRNA* flies and performed a PCR to detect the expected genomic deletion. Expected genomic deletions were identified in DNA obtained from fly heads but not in DNA obtained from the fly body suggesting tissue (eye) specific gene deletion using our newly generated reagents (Figure 4A). Further, this suggested that the *UAS-Cas9-T2A-eGFP* was indeed under the control of *eyGal4* and had minimal or no influence from other genetic elements despite being generated by random P-element insertion.

**Figure 4:**
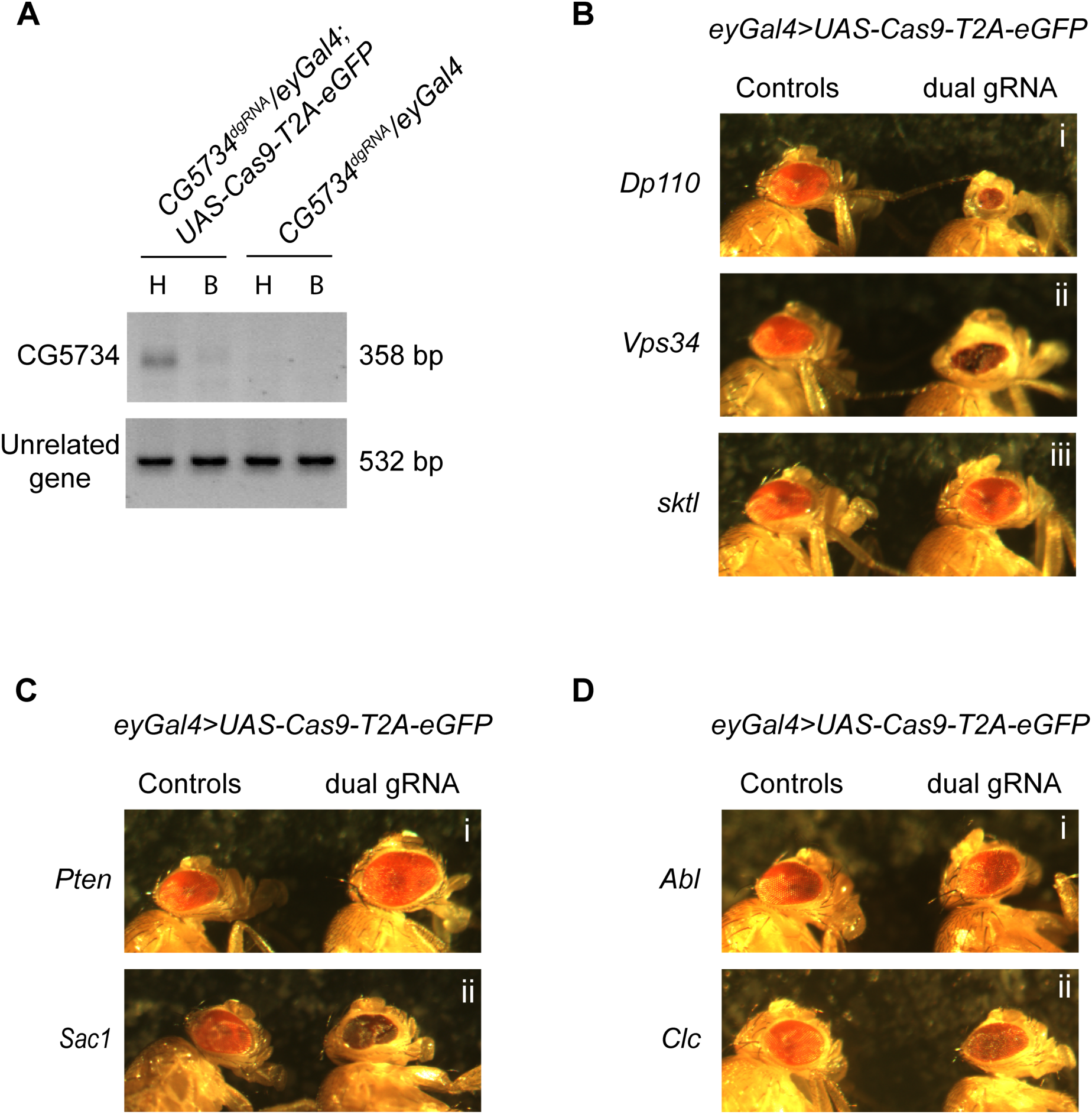
A genetic screen employing the dgRNA transgenic fly library to identify PI-signalling genes in eye development. (A) Genomic PCR of CG5734 as a representative example to show that tissue specific gene deletion can be obtained by driving UAS-Cas9-T2A-eGFP specifically in the eyes using eyGal4 in the presence of ubiquitously expressing dgRNAs. An amplicon corresponding to CG5734 deletion was obtained in DNA extracted from fly heads (‘H’) of the appropriate genotype indicated but not in DNA extracted from the bodies (‘B’) of the fly. (B) Representative images of phenotypes observed upon eye specific CRISPR mediated deletion of a few PI kinases – *Dp110, Vps34* and *sktl*. (C) Representative images of phenotypes observed upon eye specific CRISPR mediated deletion of two PI phosphatases – *Pten* and *Sac1*. (D) Representative images of phenotypes observed upon eye specific CRISPR mediated deletion of PI binding proteins *Abl* and *Clc*.

The eyes of progeny flies from each of the crosses between *ey-Gal4; UAS-Cas9-T2A-eGFP* and the 103 *U6-dgRNA* transgenic lines were imaged with appropriate controls. A spectrum of phenotypes was obtained ranging from smaller eyes as in the case of *Dp110* [Figure 4B (i)], presence of necrotic patches like with *Vps34* [Figure 4B (ii)] and larger eyes when *Pten* was knocked out [Figure 4C (i)]. *Sac1* knock out led to smaller eyes that were crumbled [Figure 4C (ii)]. CRISPR mediated knockout of PI binding proteins *Abl* [Figure 4D (i)] and *Clc* [Figure 4D (ii)] resulted in an irregular arrangement of ommatidia giving a rough appearance to the eyes. This suggested possible different roles for these proteins during eye development. In some cases, e.g., *skittles* [Figure 4B (iii)], eye morphology was comparable to the control flies with no visible morphological differences. The details of all the 103 genes and phenotypes obtained have been listed in Table I.

Deletion of Class I PI3K (*Dp110*), that converts PI(4,5)P_2_ to PI(3,4,5)P_3_ resulted in smaller eyes compared to controls. This is in agreement with a number of studies that have established the importance of PI3K for cell growth (Goberdhan et al., 1999; Weinkove et al., 1999). In order to compare the efficiency of using CRISPR based knockout versus RNAi mediated knockdown when studying tissue specific phenotypes, we expressed the PI3K dgRNA and two different PI3K^RNAi^ lines in the *Drosophila* eye using eyGal4. All three of these resulted in reduced size of the eye. While the RNAi mediated knockdown resulted in ∼40% reduction [Figure 5A (ii) and (iii) and C] compared to controls [Figure 5A (i)], dgRNA mediated PI3K knockout showed a stronger phenotype with eyes ∼75% smaller [Figure 5B (ii) and C] than appropriate controls [Figure 5B (i)] thus demonstrating that these dgRNAs serve as efficient tools to study tissue specific roles of genes.

**Figure 5:**
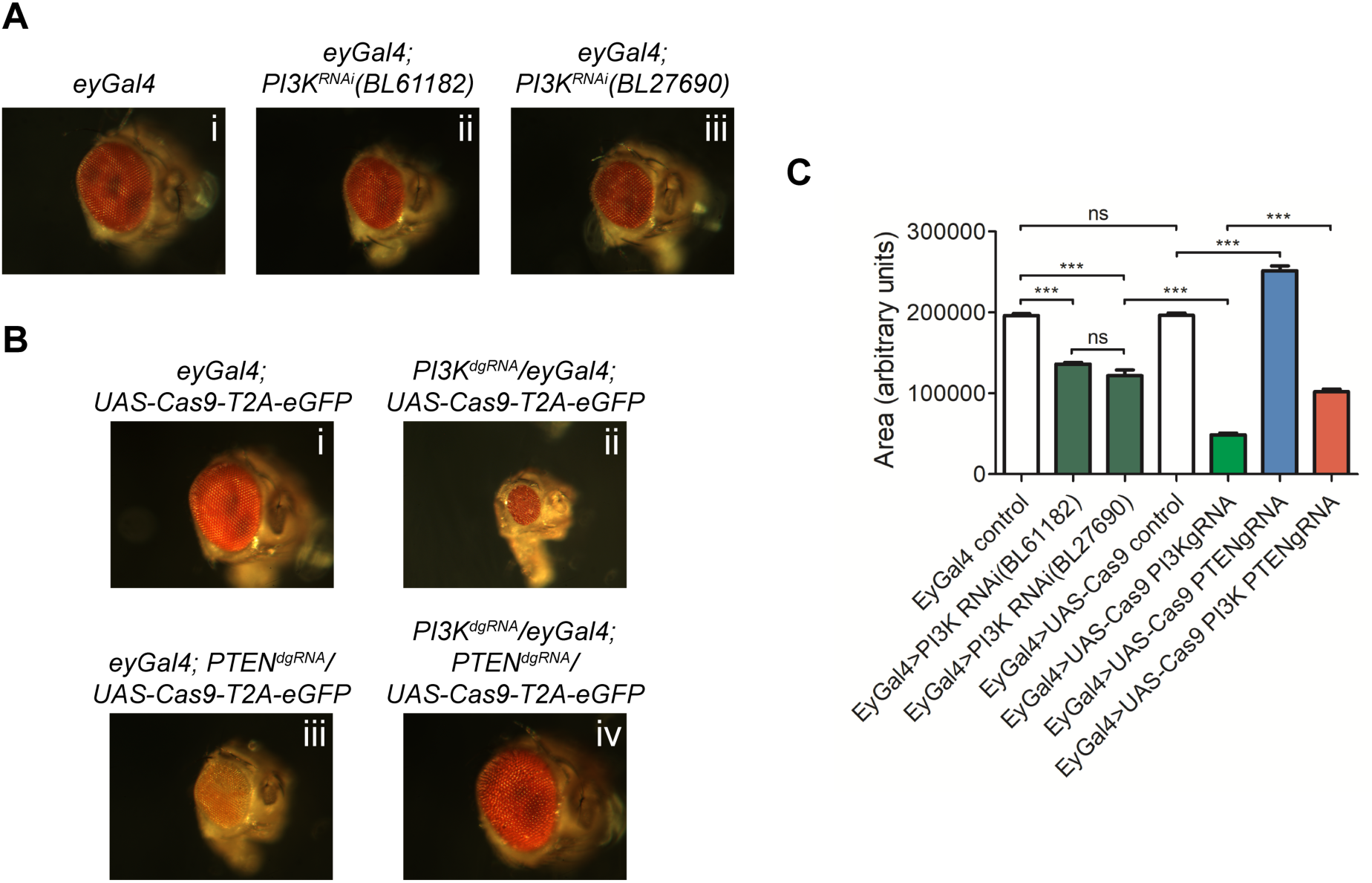
Combinatorial use of dgRNAs to study genetic interactions regulating developmental processes. (A) Knockdown of *Dp110* results in smaller eyes compared to control eyes as seen with two different RNAi lines. (B) CRISPR mediated *Dp110* knockout phenocopies RNAi mediated *Dp110* knockdown and results in smaller eyes compared to control eyes. However, the phenotype observed with the use of CRISPR to target *Dp110* is more severe compared to the phenotype seen with either of the RNAi lines. CRISPR mediated *Pten* deletion results in eyes larger than control eyes. When both *Dp110* and *Pten* are targeted simultaneously, the size of the eyes are intermediate between the small eyes seen with *Dp110* deletion and the large eyes seen with *Pten* deletion. (C) Quantifications of the phenotypes shown in Figure 4A and 4B.

### Targeting multiple genes using dgRNAs to study genetic interaction in specific tissues

Understanding genetic interactions is an important approach to establish the molecular pathways underpinning any given biological process. In order to demonstrate that the dgRNA library can be used to target multiple genes simultaneously in a tissue specific manner, we took advantage of the reversible nature of the PIP_2_ to PIP_3_ conversion by enzymes *Dp110* and *Pten*. Class I PI3K (encoded by *Dp110*) phosphorylates PIP_2_ to generate PIP_3_ while the lipid phosphatase PTEN (encoded by *Pten*) dephosphorylates PIP_3_ to generate PIP_2_. We generated a strain containing both *Dp110* dgRNA and *PTEN* dgRNA and crossed it to *ey-Gal4; UAS-Cas9-T2A-eGFP* flies. Disruption of *Pten* was able to rescue the phenotype of the disruption of *PI3K* and vice versa. The eyes of the *PI3K, Pten* double knockout flies were intermediate in size [Figure 5A (iii) and C] compared to the small eyes of the *PI3K* knockout [Figure 5A (ii)] and the large eyes of the *Pten* knockout flies (Figure 5A (iv) and C). However, the eyes of this double mutant flies were smaller compared to control flies possibly because the *Dp110* dgRNA was more efficient than the *Pten* dgRNA resulting in a greater reduction of PIP_3_ levels.

The fact that multiple genes can be targeted simultaneously in a tissue specific manner using CRISPR/Cas9 technology is of great advantage. Traditionally, genetic interactions have been studied by generation of double mutants through meiotic recombination. However, this may not be a viable option in many cases wherein the genes being studied lead to organismal lethality when disrupted or when the genetic loci of the two genes being studied are very close thereby drastically reducing the recombination efficiency between these genes. In fact, several genes involved in the PI signalling cascade are clustered together on different chromosomes. Our dgRNA transgenic fly library offers an opportunity to generate tissue specific double knockouts independent of the proximity of these genes.

### Generation of whole fly knockouts and using dgRNA transgenics

RDGB (retinal degeneration B) is a PI transfer protein (encoded by *rdgB*), depletion of which leads to light dependent phototransduction defects (Yadav et al., 2015). Whole body mutants of *rdgB* are viable with no morphological defects in the eye (Yadav et al., 2015) although electrical recordings (ERG) from the eye of *rdgB* mutants show reduced amplitude. In order to rule out the possibility that the *rdgB U6-dgRNA* was non-functional in developing eye discs, we crossed the same *rdgB U6-dgRNA* transgenic flies to *nanos-Cas9* to generate a complete germline knockout of *rdgB* (Figure 6A). ERG recordings from this germline knockout of *rdgB* generated using CRISPR/Cas9 was comparable to *rdgB*^*2*^ (Figure 6B), an amorphic allele generated by chemical mutagenesis in the Benzer lab (Harris and Stark, 1977). In a similar manner, using the *U6-dgRNA* transgenic flies, we generated germline knockouts of several additional genes by crossing them to either *nanos, vasa or Act5c-Cas9* flies (Table I). Several gRNAs when crossed to *Act5c-Cas9* or *vasa-Cas9* flies showed pupal lethality and hence were then crossed to *nanos-Cas9* flies. The deletions obtained in this manner are highly efficient (between 4-100% efficiency in F_1_ flies and between 3-70% efficiency in F_2_ flies) in generating germline knockouts with deletions ranging in size between 400bp and 38kb. For a few genes, we were unable to generate germline deletions despite repeated attempts possibly due to their essential roles in gametogenesis or embryogenesis.

**Figure 6:**
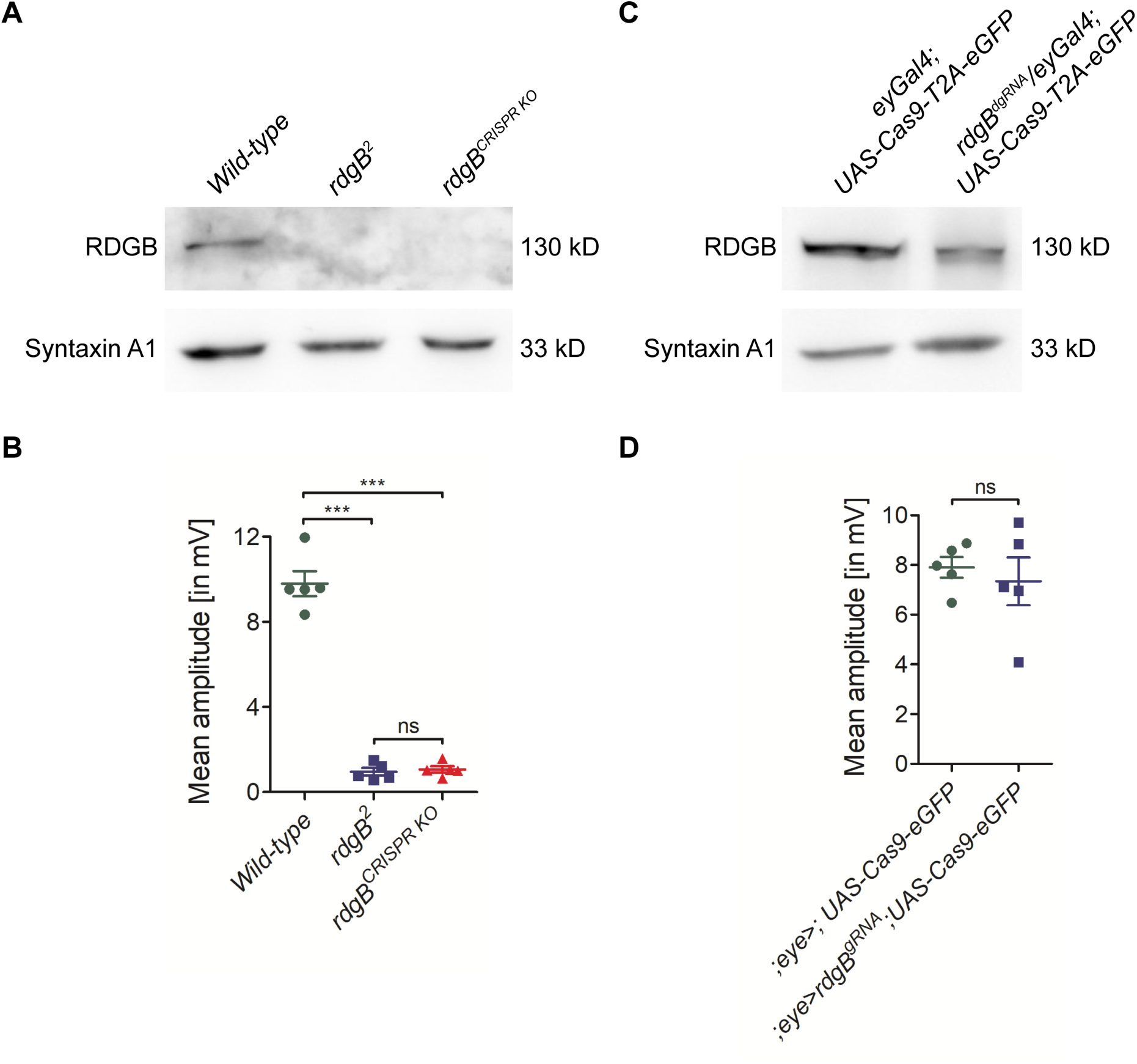
Generating germ line fly knockouts using dgRNA transgenics and anomalous behaviour of the Cas9 deletion system. The PI transfer protein RDGB (retinal degeneration B) is a good example to demonstrate both the need for whole fly knockouts and the anomalous behaviour of the Cas9 deletion system. (A) Western blot analysis showed a complete loss of RDGB in the whole fly *rdgB* deletion (*rdgB*^*CRISPR KO*^) flies, similar to the *rdgB*^*2*^ null mutant flies. (B) Both *rdgB*^*2*^ and *rdgB*^*CRISPRKO*^ flies showed complete loss of ERG amplitudes. (C) Western blot analysis showed that eye specific Cas9 expression was not sufficient to knockout *rdgB* and detectable levels of RDGB protein was still present. (D) The mean amplitude from ERG recordings of these flies was the same as control flies.

### Anomalous behaviour of the Cas9 deletion system

We crossed the *rdgB U6-dgRNA* flies with *ey-Gal4;UAS-Cas9-T2A-eGFP* flies. The progeny were collected and their ERG recorded. In contrast to the *rdgB*^*2*^ allele and the germline *rdgB* knockout generated in this study (see previous section), we observed no reduction in ERG amplitude compared to wild-type flies (Figure 6D). Although PCR analysis of these eyes confirmed the occurrence of deletion events, Western blot analysis from heads of *rdgB U6-dgRNA, ey-Gal4;UAS-Cas9-T2A-eGFP* flies revealed that the RDGB protein was still present in these eyes (Figure 6C) thus explaining the normal ERG amplitude observed. This finding implies that although *rdgB U6-*dgRNA worked efficiently to generate germline deletions, it was not effective in tissue specific depletion of RDGB protein levels. The reasons for this are not entirely clear. We speculate that RDGB is a protein with a long half-life and therefore despite efficient targeting of *rdgB* locus by the dgRNA in the eyes, the protein levels are not affected. However, it is also possible that the deletion of *rdgB* was not homogeneous in all eye precursor cells where Cas9 was expressed using *ey-Gal4*; residual photoreceptors with normal rdgB might be sufficient to generate a normal light response.

*skittles* (*sktl*) is a PI4P 5-kinase known to have a role during larval development (Hassan et al., 1998). Mutations in *sktl* are organismal lethal as well as cell lethal. Hence, we expected a strong phenotype when the *sktl* locus was deleted in eye discs using *ey-Gal4*. However, when *U6-sgRNA or U6-dgRNA* targeting *sktl* was used, we did not see any morphological phenotype in the eye [Figure 4B (iii)]. One possible reason for this could be that the *sktl gRNAs* was non-functional in eye discs although they had worked well, being able to delete *sktl* in S2R+ cells. In order to test this, we crossed *sktl U6-sgRNA* to *nanos-Cas9* flies. From this cross, heterozygous F2 progeny were collected and their *skittles* genomic locus sequenced. Out of the 141 F2 flies sequenced, 132 showed in-frame indels with an intact kinase domain, with the largest deletion being 24 bp. This suggests that the *sktl U6-sgRNA* were highly efficient at targeting *sktl* but only cells with at least a partially functional protein managed to survive during eye development and therefore appeared morphologically normal.

## Discussion

Patterning tissue architecture during metazoan development or the systemic control of animal physiology are complex processes and typically involves the function of genes acting in both cell-autonomous and non-cell autonomous modes. Identifying novel genes regulating these processes and uncovering their mode of action is facilitated by the ability to inactivate genes in specific cell types, tissue domains or organs with spatial and temporal precision. Such controlled inactivation can be achieved through the use of gene manipulation systems expressed with spatial and temporal precision using the GAL4/UAS system (Brand and Perrimon, 1993) and this approach has been coupled previously with methods to deplete specific RNAs to study their role in development and physiology (Reim et al., 2014). In this study, we present the use of the GAL4/UAS module for gene inactivation by using the CRISPR/Cas9 gene editing system. By expressing Cas9 with spatial and temporal precision using the GAL4/UAS system we were able to selectively inactivate genes in early precursor cells of the eye imaginal disc and thus uncover functions of the PI signalling system in the growth and patterning of the *Drosophila* eye. Using this method, we uncovered the function of 30 PI signalling genes in eye development and 9 genes previously not implicated in this process. Using our system, we identified phenotypes for genes such as *Dp110* and *Pten* that are previously described using either classical mutant alleles or using RNAi mediated depletion. However, the tool kit presented here includes reagents for editing ca. 72 PI binding proteins. While these have been described by *in vitro* protein biochemistry studies or predicted from structural bioinformatics, their function *in vivo* and their role in cell and developmental biology remains to be explored. Indeed, this resource can be used in almost any tissue or cell type in *Drosophila* that can be chosen for analysis, perhaps on the basis of their expression pattern or other relevant criteria. Other groups have also recognized the potential of CRISPR based mutagenesis to perform tissue specific screens in *Drosophila* and are generating gRNA transgenic fly libraries. While some studies have focussed on transcription factors, protein kinases, protein phosphatases and genes implicated in human pathologies (Port et al., 2019), others have embarked on generating large scale libraries with no special focus on any particular group of genes (Meltzer et al., 2019). Our study however represents the first comprehensive resource of editing tools that will allow a systems level analysis of PI signalling, a pathway of fundamental importance for cellular organization, tissue development and architecture.

The system we have developed includes a number of innovative features. In the study of cell-cell interactions in development, it is desirable and indeed essential to mark the cell types in which gene manipulation is being performed. To facilitate this, we have designed an *UAS-Cas9-eGFP* construct which provides many advantages: (i) All cells expressing Cas9 also express eGFP thus marking cells guaranteed to express Cas9. The expression of eGFP from this construct is a definite readout of Cas9 expression unlike the expression of eGFP from an independent UAS-eGFP transgene. (ii) Since Cas9 and eGFP are expressed as independent proteins, the possibility of a Cas9 fusion protein with reduced function is eliminated. (iii) The UAS-Cas9-eGFP presented here avoids the need for two independent genetic elements to express both Cas9 and eGFP thus reducing the complexity of strain construction. (iv) The use of Cas9 with an NLS ensures that the protein is nuclear localized maximizing the probability of gene editing activity. Thus, our system offers the ability to generate groups of gene-edited cells/tissue with high-efficiency that are guaranteed to be marked by eGFP expression.

The genetic strategy for CRISPR/Cas9 editing presented here enhances the number of genetic elements that can be brought together in a single fly and allow for CRISPR based reagents to be used in innovative ways. For example, in addition to the basic eye development screen presented here as an illustrative example, Cas9 can be expressed in a mosaic fashion under the control of the Gal4/UAS system using the CoinFLP method (Bosch et al., 2015). This would allow for the generation of CRISPR edited mosaic clones useful for studying cell-cell interactions including cell competition that are integral elements of tissue development. Indeed, a recent study has presented evidence of a role of some genes involved in PI signalling as regulators of cell competition during eye development (Janardan et al., 2019) and the expanded repertoire of editing reagents generated during this study will allow the mechanism by which PI signalling regulates cell competition to be analysed in greater detail.

Recent studies have used the approach of having two sgRNAs both targeting the coding region immediately downstream of the start codon with the expectation of maximizing the likelihood of a frameshift mutation to achieve loss of function (Port et al., 2019). By contrast, we have taken a slightly different approach of using two sgRNAs, one located near the start codon and the other near the stop codon thus generating a deletion of the full open reading frame post-editing. Even if only the gRNA targeting the first exon is functional, indels can be introduced and lead to loss of function from a resulting frameshift mutation. However, the presence of second sgRNA targeting the last exon will ensure that the entire gene is deleted thus increasing the chances of generating loss of function mutants. Secondly, because our dgRNA containing transgenic flies target both the first and the last exon and result in complete deletion of the gene, the subsequent introduction of a copy of the gene with homologous arms (Baena-Lopez et al., 2013) would result in gene knock-ins and reconstitution of function. This would require a germ line knockout of the gene and the use of our reagent set to generate such a null allele is described in the study. The use of a suitably mutagenized copy of the wild type gene would allow individual point mutations or domain deletions of a protein to be generated, for example allowing the analysis of the role of specific amino acid residues or domains in a specific cell biological question. Since such knock-in constructs would also be expressed from the same locus as the wild-type gene thereby avoiding artifacts associated with altered levels of expression when transgenes are expressed from alternate genomic loci.

During the course of this study, we identified a few limitations for the use of these reagents. For example, in the case of the PI4P 5-kinase *sktl*, eye specific deletion resulted in morphologically normal looking eyes. The reason for this was most likely that cells lacking *sktl* were eliminated (due to the essential nature of the gene product) early in eye disc development and only those cells with indels that are multiples of three bases thus possibly producing a partially functional protein survived. This is however dependent on the nature of the gene rather than the design of the gRNA constructs. For example, when sgRNA targeting the kinase domain of *sktl* was used in conjunction with *ey*>*UAS-Cas9-GFP*, flies were pupal lethal presumably because even 1-2 bp deletions within the kinase domain of *sktl* abolishes kinase activity and gene function. In the case of the PI transfer protein rdgB, we were able to detect the presence of full-length protein in flies when eye specific knockout was performed. The possibility that either of the gRNAs were inefficient or non-functional in tissues was ruled out as we were able to use the *rdgB U6-dgRNA flies* to generate a germ line knockout of this gene that phenocopied existing classical alleles. Since *rdgB* is located on the X-chromosome the inability to lose the protein by tissue specific editing is unlikely to be due to editing of only one copy of the gene. We therefore speculate that the residual protein in eye specific editing of *rdgB* most likely results from the long half-life of the protein that was present from early development. The implications of this is that when used for performing a screen in a tissue specific manner, the lack of a phenotype does not conclusively rule out the involvement of that gene. Apart from the two examples described here, it is possible that CRISPR based targeting of genes in a tissue specific manner may lead to heterogeneous disruption of the target gene especially in polyploid larval tissues such as the salivary glands and fat body. Therefore, absence of phenotypes in these tissue specific gene disruptions using dgRNA will not be conclusive and under these circumstances generation of a germ line knock out would be valuable.

In summary, we have generated a toolkit consisting of a transgenic library of 103 dgRNA and a pUAST-Cas9-T2A-eGFP that can be used in conjunction with the existing large repertoire of GAL4 lines to perform a systems level analysis of PI signalling selectively in any cell type/tissue of choice. The pUAST-Cas9-T2A-eGFP construct would be helpful to track Cas9 expressing cells in culture or in fly tissues. The U6-dgRNAs can also be transfected to delete gene function in cultured *Drosophila* cells. They can also be used to generate whole body knockouts and the null alleles so generated can be used as a template to knock in specific versions of the gene at the endogenous locus. While the PI kinases and phosphatases have been the subject of extensive analysis, the effectors of PI signalling, namely the binding proteins remain poorly studied. We envisage the availability of this dgRNA reagent set will accelerate the studies of the function of these proteins in cellular organization and tissue architecture. They will also facilitate the functional analysis of disease mechanisms in the case of those proteins linked to human disease.

## Acknowledgements

This work was supported by the Department of Biotechnology, Government of India (BT/PRJ3748/GET/l 19/27/2015), a Wellcome-DBT India Alliance Senior Fellowship (IA/S/14/2/501540) to PR and the National Centre for Biological Sciences-TIFR. We thank the *Drosophila*, Genomics and Imaging Core Facilities for extensive support in implementing this project.

